# Phylogenetic and functional diversity among *Drosophila*-associated metagenome-assembled genomes

**DOI:** 10.1101/2024.12.19.629488

**Authors:** Aaron A. Comeault, Alberto H. Orta, David Fidler, Tobias Nunn, Amy R. Ellison, Tayte A. Anspach, Daniel R. Matute

## Abstract

Host-associated microbial communities can mediate interactions between their hosts and biotic and abiotic environments. While much work has been done to document how microbiomes vary across species and environments, much less is known about the functional consequences of this variation. Here, we test for functional variation among drosophilid-associated bacteria by conducting Oxford Nanopore long-read sequencing and generating metagenome-assembled genomes (MAGs) from six species of drosophilid fly collected in association with ‘anthropogenic’ environments in North America, Europe, and Africa. Using phylogenetic analyses, we find that drosophilid flies harbor a diverse microbiome that includes core members closely related to the genera *Gilliamella*, *Orbus*, *Entomomonas*, *Dysgonomonas*, and others. Comparisons with publicly available bacterial genomes show that many of these genera are associated with phylogenetically diverse insect gut microbiomes. Using functional annotations and predicted secondary metabolite biosynthetic gene clusters, we show that MAGs belonging to different bacterial orders and genera vary in gene content and predicted functions including metabolic capacity and how they respond to environmental stressors. Our results provide evidence that wild drosophilid flies harbor phylogenetically and functionally diverse microbial communities. These findings highlight a need to quantify the abundance and function of insect-associated bacteria from the genera *Gilliamella*, *Orbus*, *Entomomonas*, and others on the performance of their insect hosts across diverse environments.

## Introduction

Host-associated microbial communities (microbiomes) affect the biology of their hosts and how those hosts interact with the environment in diverse ways. Despite the clear effects that microorganisms can have on performance and fitness of their hosts, many studies of wild populations of non-model microbiomes have characterized variation in the microbiome using 16S rRNA gene community profiling (1). While 16S rRNA gene community profiling has provided important insight into how biotic and abiotic variation can affect microbiome composition (2), we know less about how specific microorganisms—across diverse hosts and environments—affect functional or performance traits of the microbiome and host species. Estimating functional traits of specific microorganism is one way that we can begin to generate a mechanistic understanding of how members of microbial communities affect host performance.

Insects—the most diverse group of animals—provide valuable ecosystem services (e.g. pollination and nutrient cycling) (3), are important agricultural pests, and vector a range of diseases, some with economic impacts of billions of US dollars per year (4). Insect-associated microorganisms can affect these processes by detoxifying environmental and dietary toxins (5), moderating vectoral capacity (6), conferring resistance to infection of their hosts (7, 8), and even contributing to successful biological invasions (9). Among insect, fruit flies from the genus *Drosophila* (particularly *Drosophila melanogaster*) have emerged as a model system used to study host-microbe interactions (10–12). Research on host-microbiome interactions in *D. melanogaster* has largely focused on the effects of interactions with bacteria from the genera *Acetobacter* and *Lactobacillus*. For example, *Acetobacter* have been shown to promote larval development when larvae are raised on deficient diets, suggesting links between bacteria and host nutrition (13); mate choice in adult *D. melanogaster* can be influenced by *Lactobacillus* acquired through diet (14); and experimental manipulation of *Acetobacter* and *Lactobacillus* together can lead to rapid evolution in *D. melanogaster*, indicating that these bacteria are a source of selection that can contribute to local adaptation (15, 16). However, the microbial communities of wild versus lab-reared *Drosophila* differ significantly, and *Acetobacter* and *Lactobacillus* can be rare in wild *Drosophila* (10, 17). Abundances of these genera have also been shown to vary across laboratory strains of 11 *Drosophila* species (18). The few studies that estimate microbial diversity in wild *Drosophila* have shown that they harbor a more diverse microbiome than laboratory populations (10, 19). Common members of wild *Drosophila* microbiomes include bacteria from the orders Bacteriodales and Pseudomonadales, and families Orbaceae, Enterobacteriaceae, and Enterococcaceae (10, 19, 20). Therefore, while laboratory experiments in *D. melanogaster* provide evidence that symbiotic or commensal bacteria can affect diverse aspects of their host’s biology, differences between the bacteria that are the focus of laboratory studies and those reported in wild *Drosophila* highlight a need to better understand assembly rules and functional roles of members of wild *Drosophila* microbiomes.

Despite significant variation among the microbiomes of *Drosophila* species and individuals, the composition of wild *Drosophila* microbiomes suggests that these communities are not randomly assembled from a common pool of microorganisms, and patterns of co-occurrence suggest negative interactions between certain bacterial taxa (19). Recent metagenomic analyses also provide correlative evidence for functional variation among the gut bacteria associated with three mushroom-feeding *Drosophila* species (20). However, we still have a poor understanding of the functional traits possessed by specific *Drosophila*-associated bacteria. Quantifying functional variation among *Drosophila*-associated bacteria would facilitate clearer hypotheses and predictions of the impacts of host-associated bacteria on host performance and fitness. It would also allow researchers to leverage the ecological and evolutionary diversity found among species of *Drosophila* (21–24) to better understand the processes and mechanisms that shape host-microbiome interactions in the wild.

The gut microbiome of honeybees (genus *Apis*) is arguably the best functionally characterized insect microbiome (25, 26). For example, functional work on *Gilliamella* and *Snodgrassella*—core members of the bee microbiome—has shown how these bacteria possess complementary metabolic pathways and genes that affect colonization of the gut (27). *Gilliamella apicola* is also capable of metabolizing toxic sugars (28), and screens for biosynthetic gene clusters possessed by *G. apis* have identified unique ribosomally synthesized, post-translationally modified peptides (RiPPs) that protect *A. mellifera* from infection by the pathogen *Melissococcus plutonius* (7). These examples illustrate how identifying core microbiome taxa, along with using ‘bottom-up’ genome annotation approaches to predict their functional capacities, can be important first-steps towards identifying genes that mediate microbe-microbe interactions and effect host fitness.

In this study we mine whole-organism long-read Oxford Nanopore Technology (ONT) sequences to study phylogenetic and functional diversity among bacteria associated with six species of drosophilid fly. Four of the six host species we study—*D. immigrans*, *D. hydei*, *D. repleta*, and *Zaprionus indianus*—are geographically widespread and found in association with “anthropogenic” environments (sometimes referred to as “human commensals”), such as agricultural fields and compost heaps. The other two species—*Z. taronus* and *Z. tsacasi*—are forest dweller found in sub-Saharan Africa. (We note that the genus *Drosophila* is paraphyletic, with *Zaprionus* nested within *Drosophila* (24), and as such, for brevity, we include the genus *Zaprionus* when referring to “*Drosophila*” samples and bacteria throughout.) For each host sample, we identified and classified bacterial sequences from ONT sequences generated from individual wild-caught flies to estimate microbiome diversity and core microbial taxa. We then generated metagenome-assembled genomes (MAGs) for each sample and classified MAGs against the Genome Taxonomy Database. Finally, we annotated high-quality MAGs and compare functional predictions among taxonomic groups. We were able to recover a diverse set of microbial reads from whole-organism sequencing and found that our *Drosophila* samples are host to a diverse microbiome, with some ‘core’ taxa being shared among species and sample locations. Phylogenomic analyses of MAGs assembled for these core taxa show that they are phylogenetically related to bacteria from the orders Enterobacterales, Pseudomonodales, and Bacteriodales that have been sequenced from other insects, including *Gilliamella* and *Frischella* bacteria that are core members of the *Apis* (honeybees) and *Bombus* (bumble bees) gut microbiome. Finally, we show that *Drosophila*-associated MAGs from different genera vary in gene content, predicted functional enrichments, and predicted ability to produce secondary metabolic products.

## Materials and methods

### Specimen collection

We sampled populations of drosophilid flies from the United Kingdom, United States of America, and the island of São Tomé (São Tomé and Príncipe) (table S1). Individual flies were attracted to banana traps and then collected within 12 hours via aspiration or sweep netting. Flies were then briefly anesthetized with FlyNap (Carolina Biological, USA), identified under a microscope, and preserved in 100% ethanol. We focused our sequencing effort on the ‘human commensal’ species *D. hydei* (N=8), *D. repleta* (N=4), *D. immigrans* (N=13), and *Zaprionus indianus* (N=4) collected from sites in the USA, UK, and São Tomé and Principe (table S1). We also sampled one individual from each of the forest specialists *Z. taronus* and *Z. tsacasi* from São Tomé and Principe.

### DNA isolation and sequencing

DNA was isolated from individual flies using a phenol chloroform protocol developed for obtaining high molecular weight DNA from drosophilid flies for sequencing on ONT sequencers (21). Prior to DNA isolation, tissues were hydrated in hydration buffer and homogenized with sterilized pestles in tissue lysis buffer (see dx.doi.org/10.17504/protocols.io.dm6gpbdn8lzp/v2). Sequence libraries were prepared from individual HMW extractions following a modified Oxford Nanopore Ligation Sequencing Kit protocol (dx.doi.org/10.17504/protocols.io.dm6gpbdn8lzp/v2) and LSK-110 kits. We sequenced each library on individual R9.4.1 flow cells run on Oxford Nanopore MinION MK1C machines and base called raw reads (in fast5 format) with `guppy` (v6.3.8), specifying the “super high accuracy” model with the option “–config dna_r9.4.1_450bps_sup.cfg”.

### Bacterial diversity across samples

To summarize bacterial diversity, we classified sequences with `Kraken 2` (v2.1.2; (29)) against Kraken’s standard database containing RefSeq archaea, bacteria, viral, plasmid, human, and UniVec_Core sequences. We then generated BIOM-format taxonomic summaries for each sample using the `kraken-biom` tool (30). Combined BIOM tables were imported and converted into a phyloseq object in R using `import_biom` and `phyloseq` functions from the *`phyloseq`* library (31). We removed non-bacterial taxa and taxa with low variance across samples (threshold = 1 x 10^-7^). To identify the most abundant bacteria within each sample we first visualized variation in the proportion of reads assigned to bacterial classes. To summarize differences in the relative abundances of bacteria across samples we converted the phyleseq object to a DGElist using the ‘`phyloseq_to_edgeR`’ function from the *`PathoStat`* library (32) and conducted a Multidimensional scaling analysis with the `plotMDS` function from the *`limma`* R package using the “pairwise” gene comparison method. Because sequencing resulted in uneven sequencing depths across samples, we conducted a redundancy analysis (RDA) using the `rda` function from the *`vegan`* R library. We specifically modeled taxonomic abundance (read counts per bacterial taxon) as a function of the interaction between total sequencing depth and host species.

### Metagenome assembly and classification

Using read-level classifications generated by `Kraken2`, we identified and isolated bacterial reads from whole-organism fastqs by selecting reads that were not unclassified or classified as "Homo" using `seqtk`’s ‘`subseq`’ command (v1.3-r106; https://github.com/lh3/seqtk). We then assembled metagenomic contigs from the pool of putatively bacterial reads with `metaFlye` (v2.9; (33)), and polished assembled contigs with `medaka` (v1.7.2; (34)). Polished contigs were binned into MAGs using `metaBAT2` (35) and levels of completeness and contamination were assessed using `CheckM` (‘lineage_wf’; v1.1.3; (36)).

We used `GTDB-TK`’s ‘`classify_wf`’ pipeline (37) to assign taxonomy and determine phylogenetic relationships among MAGs with completeness greater than 45% and contamination less than 10% (as determined by `CheckM`). `GTDB-TK` leverages the Genome Taxonomy Database (38) and tools that allow for taxonomic assignment through sequence clustering, alignment, and large-scale phylogenetic reconstruction (39–44). To explore relationships among drosophilid MAGs and publicly available bacterial genomes, we also carried out focused analyses on MAGs from bacterial genera that were abundant across our set of assembled MAGs (see results) using `GTDB-TK`’s ‘`de_novo_wf`’ pipeline. Similar to `classify_wf`, the `de_novo_wf` pipeline uses `Prodigal` (40) and `HMMER` (39) to identify marker genes in each MAG. Genes are then concatenated and phylogenetic relationships are inferred using `FastTree` (43) run with the WAG+GAMMA model. These analyses applied taxonomic filters at the order level and required a minimum of 50% of amino acids for a given MAG to be included in the alignment for that MAG to be retained in the analysis. We subsequently implemented `de_novo_wf` analyses separately for the orders Enterobacterales, Bacteriodales, and Pseudomonodales (outgroups: Pseudomonadales, Sphingobacteriales, and Enterobacterales, respectively). Finally, we manually pruned phylogenetic trees produced by this analysis and compared the host organisms of publicly available bacterial genomes that were within the same genera as focal drosophilid MAGs.

### Annotation and functional characterization of MAGs

We annotated MAGs that received `CheckM` completeness and contamination scores greater than 45% and 10%, respectively, using `eggNOG-mapper` v2 (45, 46) run with `eggNOG v5.0` (47) and protein predictions made by `Prodigal` (40). We focused functional comparisons on lineages of bacteria for which we had multiple MAGs assembled from either different drosophilid species or different locations within the same species. Specifically, we focused on MAGs assembled for bacterial taxa within the families Enterobacterales (N=27 MAGs), Pseudomonodales (N=21), and Bacteroidales (N=14), as species within these families are likely to be core (or symbiotic) members of the microbiomes of the *Drosophila* species we sampled (see Results). For gene sets annotated for each MAG within these three families, we identified COG categories and KEGG Orthology (KO) pathway maps that were enriched at the genome level using COG and KEGG annotations generated by `eggNOG-mapper` and the ‘`enrichCOG`’ and ‘`enrichKO`’ functions in the *`MicrobiomeProfiler`* R library (v1.6.1; (48)).

We compared functional diversity among groups of *Drosophila*-associated bacteria by identifying COG categories and KEGG pathway maps that were enriched in at least 80% of the focal group being assessed and in less than 50% of all other MAGs in our dataset. We tested for functional enrichment among MAGs from each of the genera *Gilliamella*, *Orbus*, *Acinetobacter*, *Entomomonas*, *Pseudomonas*, and *Dysgonomonas*. We also compared enrichment in *Gilliamella* and *Orbus* MAGs we assembled as part of this study with enrichment in 12 publicly available *Gilliamella* genomes derived from *Apis* bees and 13 derived from *Bombus* bees (table S3; *Gilliamella* clade in fig 4A).

Finally, we predicted biosynthetic gene products produced by MAGs classified in the orders Enterobacterales, Pseudomonodales, or Bacteroidales using the ‘antibiotics and secondary metabolite analysis shell’ (`antiSMASH` v7; (49)). We first annotated genes in each MAG belonging to these three families using `Prokka` (v1.14.5) run with default parameters. `Prokka` annotations were then used as input to antiSMASH. We ran the online version of antiSMASH (https://antismash.secondarymetabolites.org/; (49)) with relaxed detection strictness and all options on. For visualization and analyses we labeled singleton biosynthetic classes as “Others” and nonribosomal peptides’ hybrid classes as “Other NRPS”. We performed Kruskal–Wallis (KW) and post-hoc Wilcoxon ranks sum test to determine differences of the frequency of BGCs per MAG between bacteria orders. We fit a linear model to test for effects of genome size and bacterial order on the (log_10_) number of BGCs per MAG. In this analysis we first tested for an interaction between genome size and bacterial order, but this was not significant; we therefore include genome size and bacterial order as predictors in the final model, but not the interaction between the two. Finally, to test whether the presence/abundance of BGCs classes varied across bacterial orders we conducted a permutational analysis of variance (PERMANOVA) on Euclidean distances between MAGs as implemented with the `adonis` function in the `vegan` package in R (Oksanen et al. 2020).

## Results

### Bacterial diversity across sampled reads

By mining whole-organism sequence reads with `Kraken2` we recovered a median of 198,036 bacterial reads (range: 30,630-5,574,666) spanning a median of 765.21 Mbps of sequence (range 197.91-8,829.46 Mbps) per individual drosophilid fly (table S1; fig. S1). Across all samples, sequence reads were uniquely assigned to 12 phyla, 23 classes, 62 orders, 114 families, and 254 genera of bacteria. Gammaproteobacteria, Alphaproteobacteria, Bacilli, and Mollicutes were each the most abundant class of bacteria in sequences obtained from 22, 4, 3, and 2 individuals, respectively; and Gammaproteobacteria, Alphaproteobacteria, and Bacilli were the second most abundant classes of bacteria in individuals where they were not the most abundant, except for one individual with Epsilonproteobacteria being the second most abundant class (13.4% of reads) and three individuals with Flavobacteria as the second most abundant class (13.1% to 26.0% of reads) (fig. 1A; fig. S2). The percentage of reads assigned to the most abundant class of bacteria per individual ranged from 25.0% to 90.2%, while the percentage of reads assigned to the second most abundant class of bacteria per individual ranged from 3.9% to 43.1% (fig. 1A). Reads assigned to *Acetobacter* and *Lactobacillus*—the two genera of bacteria that are the focus of laboratory research in *Drosophila melanogaster*—were relatively rare across our samples, with median proportions of reads per sample of 0.18% (90% empirical quantile: 0.03% to 8.25%) and 0.22% (0.04% to 2.51%), respectively. Inspecting rarefaction curves showed that taxonomic classification across reads was saturated for ∼50% of sampled *Drosophila*; however, taxon sampling was not saturated in individuals where we recovered fewer than ∼100,000 bacterial reads (fig. S3; table S1).

**Figure 1.**
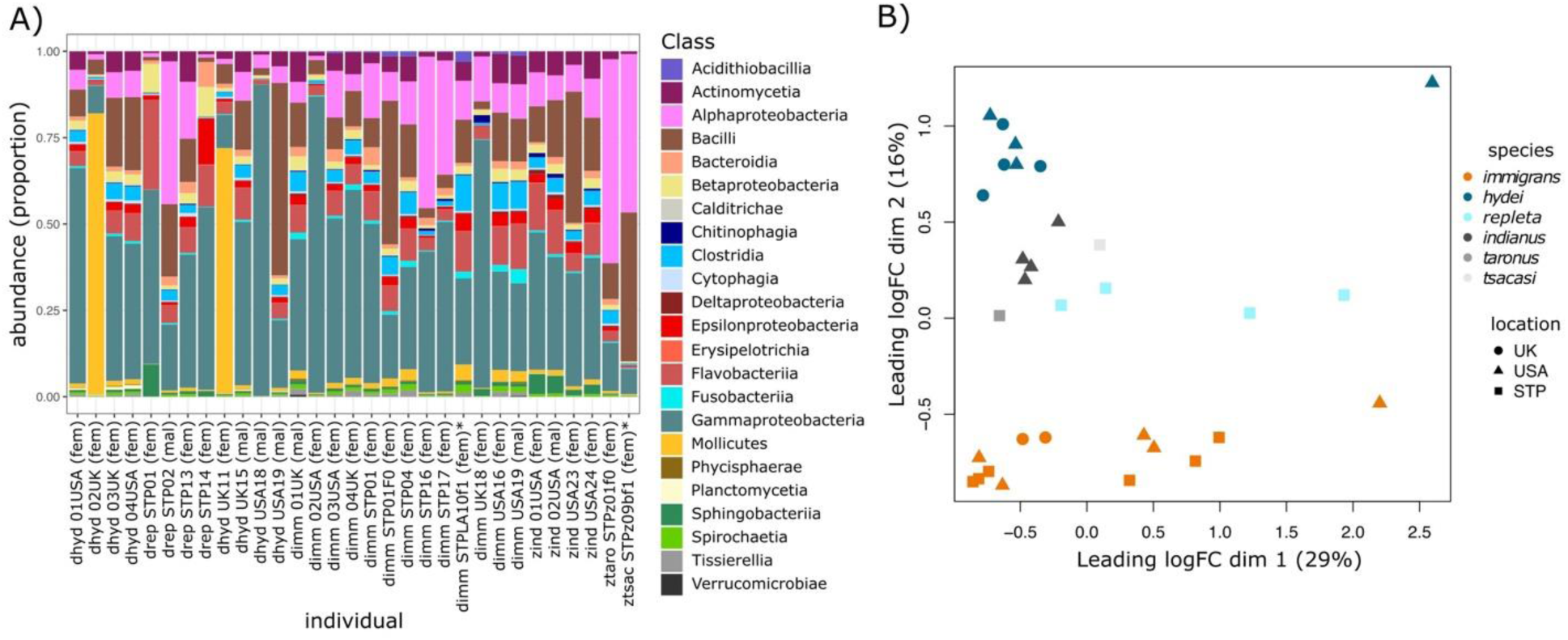
Bacterial sequence reads derived from whole-organism extraction and Nanopore Minion sequencing represent diverse taxa belonging to 23 classes (**A**). Differences in the relative abundances of microbial taxa among samples were affected by the number of sequences classified as bacterial (**B**; dim 1) and by species-level differences in the microbial community (**B**; dim 2). In panel A, individual names include details of the species (dhyd = *D. hydei*; drep = *D. repleta*; dimm = *D. immigrans*, zind = *Z. indianus*, ztaro = *Z. taronus*, ztsac = *Z. tsacasii*), location (USA, UK, or STP [São Tomé and Principe]), and sex (fem = female, mal = male). Two individuals in the data set were F1 offspring from wild-caught females and are indicated with * in panel A.

We summarized differences in microbial taxa abundances among samples by first conducting a multidimensional scaling analysis. The first principal coordinate axis (PCoA) explained 29% of variation in leading log2-fold changes in bacterial read abundances among samples (fig. 1B); however, PCoA dimension 1 scores are correlated with the number of bacterial sequences recovered from a sample (Pearson’s rho = 0.63; *P* < 0.00015). The second PCoA accounted for 16% of variation among samples and differentiated samples based on host species rather than the location the hosts were collected from (fig. 1B). Redundancy analysis on the taxonomic matrix revealed that the interaction between sequencing depth and host species significantly affected bacterial diversity and abundance (model R^2^ = 81.35%; permutation test: *F*_9,21_ = 15.54; *P* = 0.003; fig. S4). Classification and analysis of bacterial sequences therefore allowed us to quickly identify diverse bacterial communities that varied among host species. However, we do not explore community diversity further because the ONT sequencing we conducted is PCR-free whole-genome shotgun sequencing, therefore variation in read abundance could be affected by factors that include differences in taxonomic abundance, genome size, and sequence classification accuracy among bacterial taxa and across their genomes.

### Bacterial diversity across MAGs

Across all samples, we assembled 143,751 contigs that were then binned into 366 MAGs (table S1). We retained 103 ‘focal’ MAGs after filtering for `CheckM` completeness scores greater than 45% and contamination less than 10%. We were unable to recover MAGs from six of our 31 sampled host individuals, and both the number of contigs and MAGs assembled from a sample were correlated with the amount of bacterial sequence data recovered from the whole-organism sequences (Kendall’s tau = 0.65 and 0.58, respectively; both *P* < 1 x 10^-5^; fig. S5).

Largely consistent with read-level classifications reported by `Kraken2`, the most abundant classes of bacteria across the 103 focal MAGs were Gammaproteobacteria (55 MAGs), Bacteroidia (21 MAGs), Bacilli (11 MAGs), Alphaproteobacteria (10 MAGs), Clostridia (4), and Camylobacteria (2). Using GTDB-TK taxonomy, we identified seven orders that were represented by at least 5 MAGs: Enterobacterales (N=27), Pseudomonodales (N=21), Bacteriodales (N=14), Lactobacillales (N=9), Acetobaterales (N=7), Flavobacteriales (N=5), and Burkholderiales (N=5) (fig. 2A). Three of these orders—Enterobacterales, Pseudomonodales, and Bacteriodales—were represented by multiple MAGs assembled from at least two host species, and within at least one of those hosts they were assembled from samples collected at multiple locations (fig. 2B-D). Within these three orders, seven genera were represented by MAGs assembled from *D. hydei*, *D. repleta*, and *D. immigrans* (our most widely sampled species) and/or from multiple geographic locations (fig. 3). We focused functional analyses on the 62 MAGs from bacteria belonging to these three orders and seven genera because they are either ecologically associated with a common environment used by the species we sampled, or they are evolutionarily associated as core members of the human- commensal drosphilid microbiome.

**Figure 2.**
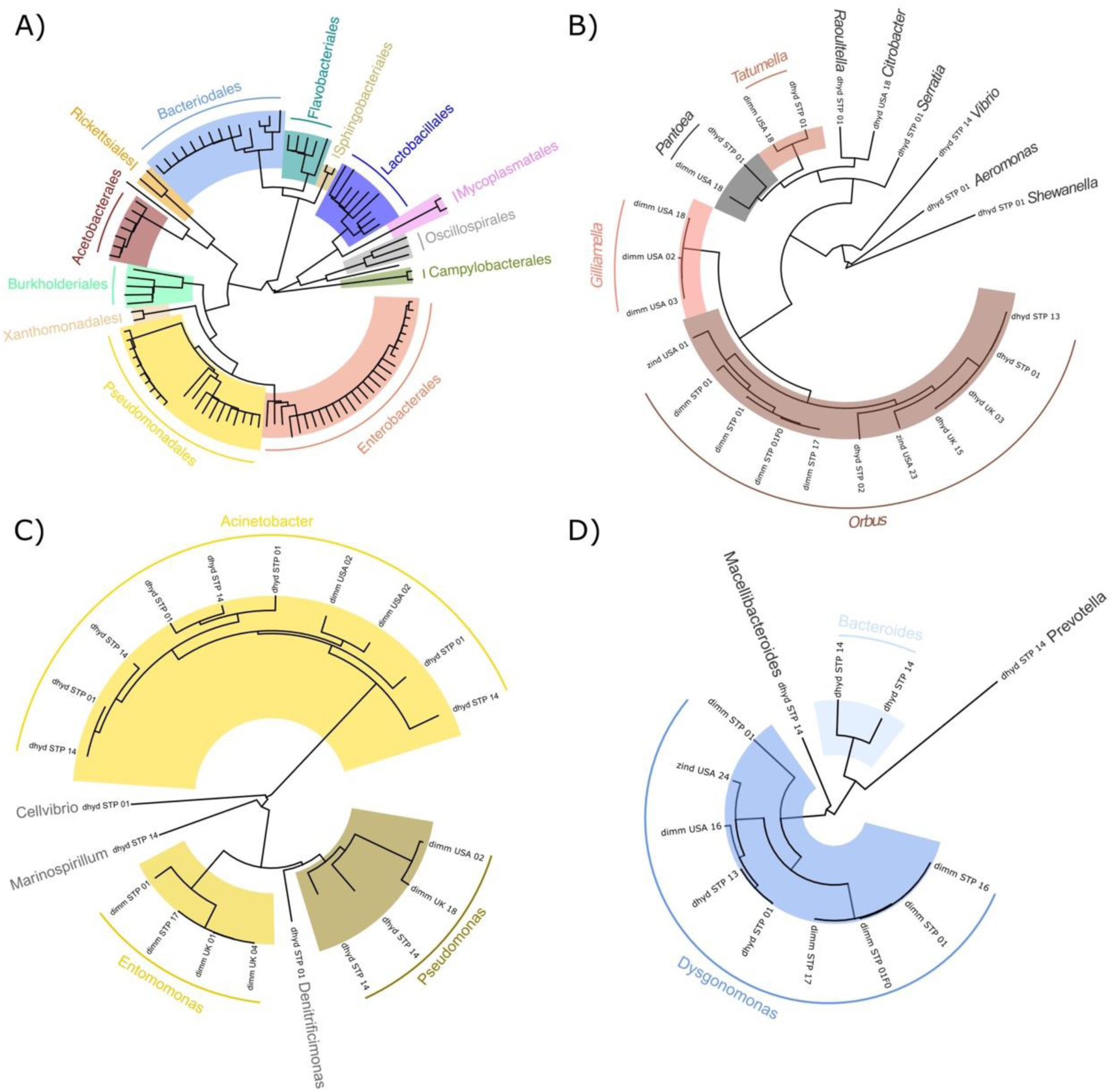
Phylogenetic classifications of drosophilid-derived MAGs. A) Phylogenetic relationships among *Drosophila* MAGs from maximum-likelihood placement by pplacer as implented in GTDB-TK’s ‘classify_wf’ pipeline against the GTDB-Tk reference tree. Phylogenetic relationships among *Drosophila* MAGs from the three most abundant orders—Enterobacterales (B), Pseudomonadales (C), and Bacteriodales (D)—generated using GTDB-Tk’s ‘de_novo_wf’ pipeline and highlighting genera represented by multiple MAGs within each order. In panels B through D, details on the host species, individual sample identifier, and location, are reported in tip names (details as in Figure 1).

**Figure 3.**
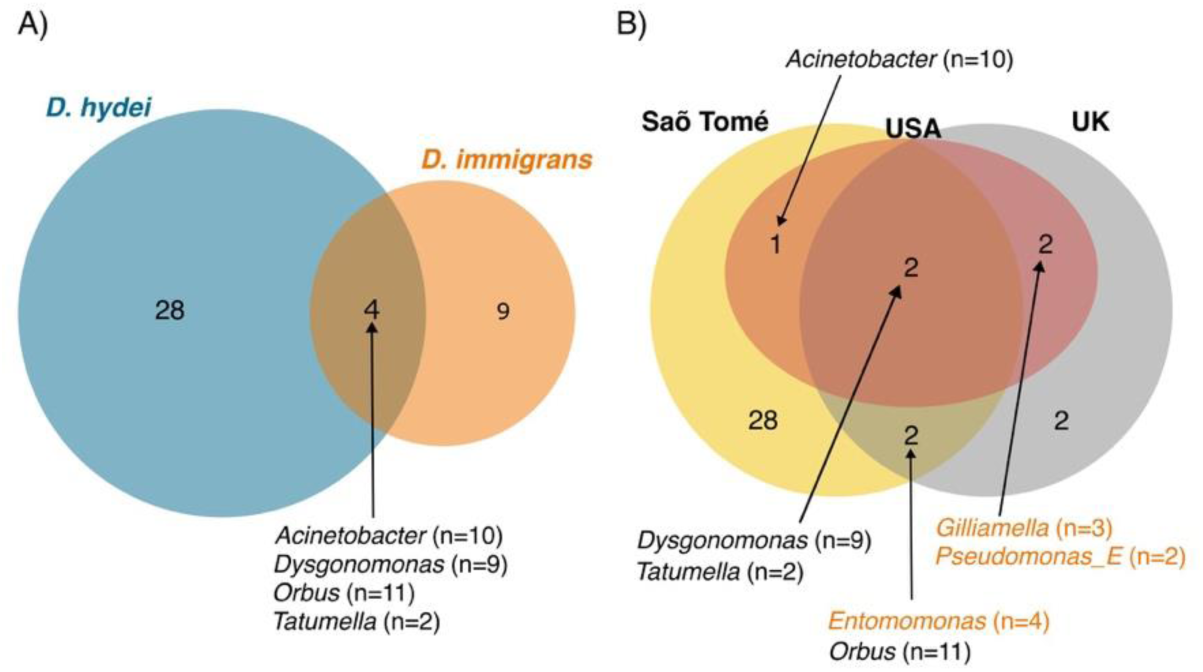
Core bacterial genera are shared among *Drosophila* species (**A**) and across geographic regions we sampled (**B**). Bacterial genera (number inside circles) and the number of MAGs assembled for each genus (beside generic names) are given for each scenario of overlap. Bacterial genera highlighted in orange (*Entomomonas*, *Gilliamella*, and *Pseudomonas_E*) were only assembled from *D. immigrans* hosts. Note that numbers of MAGs belonging to *Dysgonomonas* and *Orbus* include MAGs assembled from *Z. indianus* (see Figure 2), and *D. repleta* were grouped with *D. hydei* for this analysis.

Endosymbiotic bacteria sequences were also abundant in four of our 31 samples: sequences classified as *Wolbachia* were abundant in *Z. taronus* and *Z. tsacasi* collected on São Tomé (39,265 and 2,675 reads, respectively), and high-quality MAGs classified in the diverse group of *Wolbachia pipientis* were assembled from both samples (CheckM completeness > 99% and contamination = 0.00%; table S2). Sequences classified as *Spiroplasma* were abundant in two *D. hydei* collected from the UK (86,369 and 76,409 reads), and MAGs classified as *Spiroplasma poulsonii* were assembled from both samples (CheckM completeness = 98.5% and 72.18% and contamination = 3.01% and 0.00%, respectively; table S2).

To explore the hypotheses that ‘focal’ bacterial genera are either found in a common environment shared among human commensal drosophilids or are evolutionarily associated members of the core drosophilid microbiome, we compared MAGs to publicly available bacterial genomes using `GTDB-TK`’s ‘`de_novo_wf`’ pipeline. Drosophilid-derived MAGs classified as *Gilliamella* and *Orbus* were found to be closely related to bacteria belonging to the genera *Gilliamella*, *Orbus*, and *Frischella* that have been assembled from diverse honey bee (*Apis*) and bumble bee (*Bombus*) hosts (fig 4A) (27, 50). *Gilliamella* and *Orbus* MAGs derived from the drosophilid hosts we sampled tended to form monophyletic clades relative to bacteria from honey or bumble bee hosts, suggesting host-specific divergence (fig 4A). Orbales have also been reported in metagenomic studies of wild mushroom-feeding *Drosophila* (19, 20), suggesting broad associations between these bacteria and drosophilid hosts. Drosophilid-derived MAGs classified in the genus *Dysgonomonas* were found to be closely related to *Dysgonomonas* genomes derived from diverse sources, including environmental samples, insects, and mammals (fig 4B; table S3); and drosophilid- derived MAGs classified in the genus *Entomomonas* were related to two *Entomomonas* species in the `GTDB-TK` database—one was derived from an eastern honey bee (*Apis cerana*) host and the other from a house cricket (*Acheta domesticus*) (fig 4C). Phylogenetic comparisons to publicly available genomes therefore suggest that many of the MAGs from the genera *Gilliamella*, *Orbus*, and *Entomomonas* are associated as core members of the insect (gut) microbiome.

**Figure 4.**
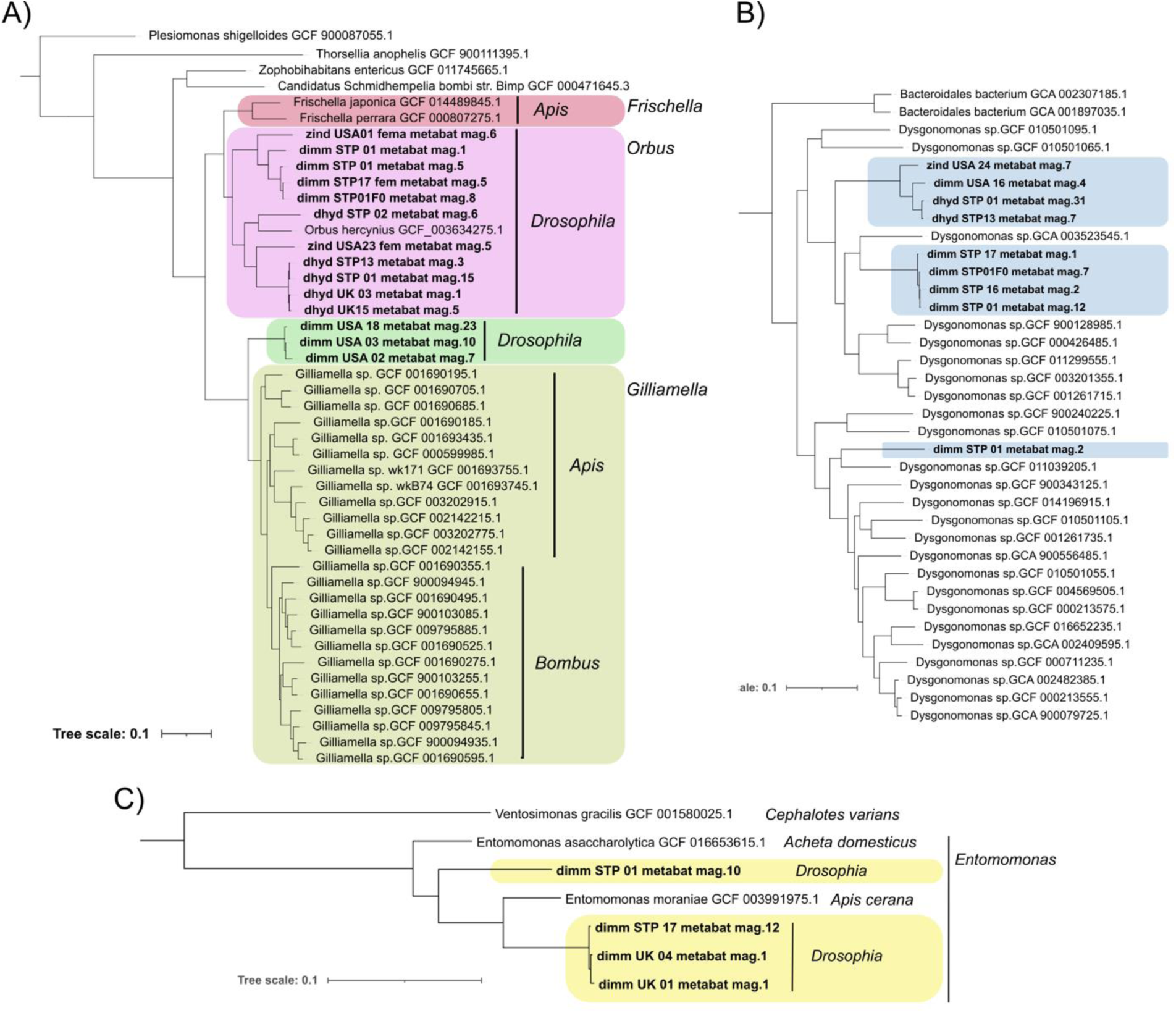
Phylogenetic relationships among focal MAGs assembled from drosophilid flies (bold type) and publicly available genomes included in GTDB-Tk’s ’de_novo_wf’ pipeline. **A**) Enterobacterale MAGs from drosophilid hosts are closely related to genomes from the genera *Gilliamella*, *Orbus*, and *Frischella* that have been assembled from diverse honey bee (*Apis*) and bumble bee (*Bombus*) hosts. **B)** Drosophilid MAGs from the genus *Dysgonomonas* are closely related to diverse *Dysgonomonas* genomes assembled from insect, mammal, and environmental sources (sources not shown in panel B). **C)** *Drosophilid* MAGs from the genus *Entomomonas* are closely related to genomes from *Entomomonas* sequenced from the eastern honey bee (*Apis cerana*) and the house cricket (*Acheta domesticus*).

### Functional variation among drosophilid-associated bacteria

We quantified enrichment in COG categories and KEGG pathways among MAGs belonging to the seven focal bacterial genera described above to test for evidence of functional differences among them. We found evidence for variation in enrichment of 12 COG categories and 27 KEGG pathways across genes annotated in each MAG (fig. 5). Comparing enrichment of KEGG pathways among bacterial genera found that greater than 80% of drosophilid-derived *Gilliamella* or *Orbus* genomes are enriched for genes belonging to 10 KEGG pathways that are under-enriched (i.e. fewer than 50% of MAGs with enrichment) in the 35 Pseudomonadales and Bacteriodales MAGs in our dataset (fig 5B). *Acinetobacter, Entomomonas,* or *Pseudomonas* genomes are enriched for genes belonging to 14 KEGG pathways that are under-enriched in the 43 Enterobacterales and Bacteriodales MAGs in our dataset (fig 5C), while *Dysgonomonas* genomes are enriched for genes belonging to 4 KEGG pathways that are under-enriched in the 48 Pseudomonadales and Enterobacterales MAGs in our dataset (fig 5D).

**Figure 5.**
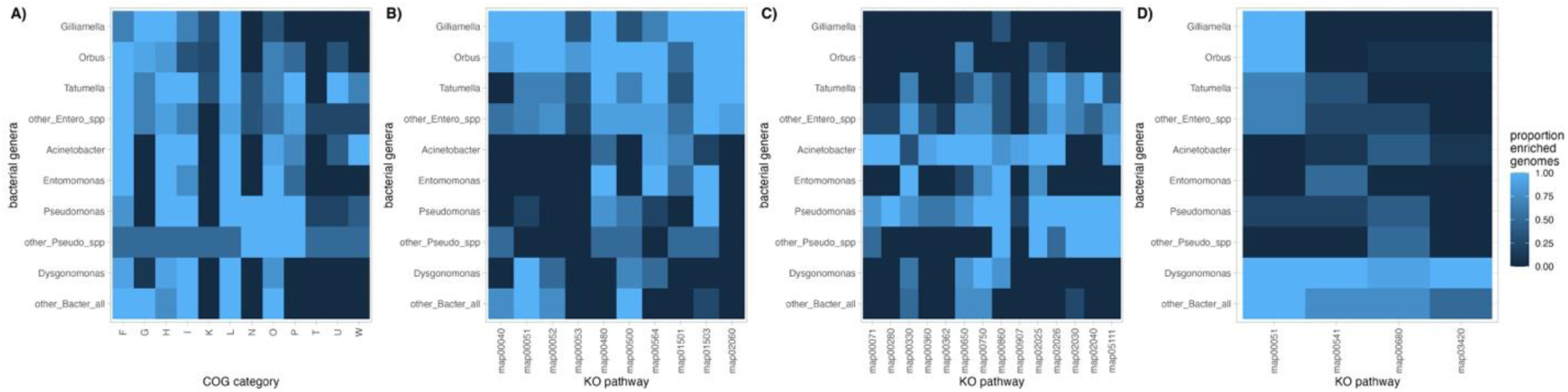
MAGs belonging to different bacterial genera differ in their functional gene content. **A)** COG categories that show variation in enrichment among MAGs. **B)** KEGG pathways that are enriched in greater than 80% of the MAGs assembled from *Gilliamella* or *Orbus* (order: Enterobacterales) but were enriched in fewer than 50% of other MAGs in our dataset. **C)** As in B), but with overrepresented enrichment within genera *Acinetobacter*, *Entomomonas*, or *Pseudomonas* (order: Pseudomonadales). **D)** As in B), but with overrepresented enrichment within the genus *Dysgonomonas* (order: Bacteriodales).

KEGG pathways that are enriched in *Gilliamella* or *Orbus* MAGs tend to be enriched across both genera: an average of 87.5% of MAGs in these genera show enrichment in the same 10 KEGG pathways (range = 58.3 to 100%; fig 4B). Seven of the 10 KEGG categories that are enriched in *Gilliamella* and *Orbus* MAGs are also enriched in more than 50% of other Enterobacterales MAGs (map00051, map00052, map00480, map00500, map00564, map01503, and map02060), suggesting functions associated with these pathways are shared across the Enterobacterales species associated with drosophilid hosts. However, three KEGG pathways (map00040, map00053, and map01501) are enriched in fewer than 50% of other Enterobacterales MAGs, suggesting possible unique or enhanced functions within *Gilliamella* and/or *Orbus*. Two of these three pathways contain genes involved in carbohydrate metabolism— specifically in pentose and glucuronate interconversions (map00040) and ascorbate and aldarate metabolism (map00053)—, while the third (map01501) is involved in resistance to beta-Lactam antibiotics. Interestingly, map00053 is only enriched in one of the three *Gilliamella* MAGs while map01501 is enriched in all three *Gilliamella* MAGs and 6 of the 12 (50%) *Orbus* MAGs, suggesting potential functional differences between strains of these closely related genera.

In contrast to Enterobacterales, there is more variation in enrichment among Pseudomonadales from the genera *Acinetobacter, Entomomonas,* and *Pseudomonas*: an average of 62.7% of MAGs from these genera show enrichment in the same KEGG pathway (range = 33.3% to 93.3%; fig 5C). Within Bacteriodales, the four KEGG pathways that are enriched across the 10 drosophilid-derived *Dysgonomonas* MAGs tend to be enriched in the other Bacteriodales MAGs (50% - 100%); however, we note that there are only 4 MAGs assigned to the order Bacteriodales that were not within the genus *Dysgonomonas* in our dataset. Nonetheless, all four KEGG pathways enriched in *Dysgonomonas* (map00051, map00541, map00680, and map03420) were rarely enriched in non-Bacteriodales (44.2%, 16.9%, 20.1%, and 2.3%, of MAGs with enrichment, respectively; fig 5D). Map00051 was enriched in *Dysgonomonas, Gilliamella,* and *Orbus* genomes (fig 5D, first column) and is involved in the metabolism of fructose and mannose sugars. The other three pathways contain genes involved in glycan sugar biosynthesis and metabolism, methane metabolism, and nucleotide excision repair. Taken together, variation in KEGG pathway enrichment across *Drosophila*-associated MAGs indicate that bacterial members of the *Drosophila* microbiome are functionally different; for example, via differences in metabolic capacity for various substrates (e.g. carbohydrates versus methane) or the ability to resist environmental stressors (e.g. antimicrobials or general DNA damage).

We also found that *Drosophila*-associated *Gilliamella* and *Orbus* MAGs differed in functional enrichment when compared to publicly available *Gilliamella* genomes isolated from *Apis* and *Bombus* hosts. COG categories N (cell motility) and U (intracellular trafficking, secretion, and vesicular transport) are enriched in *Apis*- and/or *Bombus*- associated genomes, but are not enriched across the majority of *Drosophila*-associated *Gilliamella* or *Orbus* MAGs (however, 4/11 [36.36%] of the *Orbus* MAGs isolated from *Drosophila* were enriched for COG category U). In addition to COG categories, 11 KEGG map pathways varied in enrichment among *Gilliamella* (*Drosophila*), *Orbus* (*Drosophila*), *Gilliamella* (*Apis*), and *Gilliamella* (*Bombus*) MAGs or genomes (table 1). Four pathways (map00053, map00561, map00561, and map00630) are enriched in *Orbus* genomes derived from *Drosophila* hosts relative to genomes from the other four groups. These pathways contain genes involved in the metabolism of various carbohydrates and lipids (table 1.). One of these pathways (map00630) is also enriched in *Gillimella* genomes derived from hosts in the genus *Apis*. Three pathways (map00261, map00450, and map01501) are enriched in *Gilliamella* derived from *Drosophila* hosts: maps 00261 and 01501 are both involved in antimicrobial production, and map01501 was also less likely to be enriched in *Drosophila*-derived MAGs that were not in the genus *Gilliamella* (fig. 5B). Four pathways involved in biofilm formation or movement (map02025, map02026, map02030, and map02040) are enriched in *Gilliamella* derived from both *Bombus* and *Apis* hosts. The two pathways involved in biofilm formation— map02025 and map02026—are also enriched in 5 (45%) and 3 (27%) of the *Orbus* MAGs, respectively. While the KEGG pathways above highlight potential functional differences among *Orbaceae* from different hosts, we also found 51 KEGG pathways that are enriched across all *Orbaceae* genomes (table S4). These pathways highlight that *Orbaceae* also share diverse metabolic and biosynthetic pathways.

**Table 1.** KEGG pathways that showed variation in enrichment across genomes from *Orbus* from *Drosophila* hosts, *Gilliamella* from *Drosophila* host, *Gilliamella* from *Apis* host, or *Gilliamella* from *Bombus* host (see fig. 4A for relationships among these groups). The proportion of genomes of each group that showed enrichment in each pathway is given under columns “*Orbus*”, “*Gill.* (*Dros*.)”, “*Gill*. (*Apis*)”, and “*Gill*. (*Bombus*)”, respectively. Values above 0.8 are highlighted in bold italic font.

| KEGG ID | pathway name | *Orbus* | *Gill. (Dros.)* | *Gill (Apis)* | *Gill (Bombus)* |
| --- | --- | --- | --- | --- | --- |
| map00053 | Ascorbate and aldarate metabolism | **0.82** | 0.33 | 0.25 | 0.08 |
| map00450 | Selenocompound metabolism | **0.82** | **1.00** | 0.50 | 0.00 |
| map00561 | Glycerolipid metabolism | **0.82** | 0.00 | 0.33 | 0.00 |
| map00564 | Glycerophospholipid metabolism | **1.00** | 0.67 | 0.08 | 0.15 |
| map00630 | Glyoxylate and dicarboxylate metabolism | **1.00** | 0.33 | **0.83** | 0.15 |
| map00261 | Monobactam biosynthesis | 0.45 | **1.00** | 0.08 | 0.00 |
| map01501 | beta-Lactam resistance | 0.55 | **1.00** | 0.17 | 0.38 |
| map02025 | Biofilm formation - *Pseudomonas aeruginosa* | 0.45 | 0.00 | 0.42 | **0.85** |
| map02026 | Biofilm formation - *Escherichia coli* | 0.27 | 0.00 | **1.00** | **1.00** |
| **map02030** | Bacterial chemotaxis | 0.00 | 0.00 | **1.00** | **1.00** |
| **map02040** | Flagellar assembly | 0.00 | 0.00 | **1.00** | **1.00** |

### Biosynthetic potential of drosophilid-associated bacteria

We used `antiSMASH` to annotate secondary metabolite biosynthetic gene clusters (BGCs) in 56 of the 62 MAGs classified as Enterobacterales, Pseudomonadales, or Bacteroidales (fig. 6A). Across MAGs we identified 177 biosynthetic gene clusters from 31 small molecule classes and their hybrids (table in S5). Across the 56 MAGs that carried at least 1 BGC, we annotated an average of 3 BGCs per MAGs, found more BGCs per MAG for MAGs from the order Pseudomonadales (maximum of 14 BGCs per MAG) (KW χ^2^ = 14.683, *p* < 0.001) (fig. S6A), and found a significant positive relationship between the genome size and the number of BGCs per MAG (ANOVA on genome size; *F* = 48.668, *P* = 3.8x10^-09^; fig. S6B). The most abundant class of BGC was post-translational modified peptides (RiPP-like) (n=40); followed by aryl polyene (n=28), nonribosomal peptides’ hybrids (Other NRPS) (n=22), nonribosomal peptides (NRPS) (n=19) and beta-lactone (n=15) (fig. 6B). In total, MAGs classified as Pseudomonadales possessed 91 biosynthetic gene clusters from 24 BGC classes, Enterobacterales possessed 68 biosynthetic gene clusters from 16 BGC classes, and Bacteroidales possessed 18 biosynthetic gene clusters from 9 BGC classes. The relative abundance of BGC classes differed among our three focal bacterial orders (Permutational MANOVA: *F* = 3.722; *P* = 0.001; fig 6): for example, 32% of BGCs in Enterobacterales were in the class RIPP-like, while 20% were RIPP-like in Pseudomonadales, and RIPP-like BGCs were absent in Bacteroidales. Aryl polyene class made up 20%, 12%, and 11% of BCGs annotated in Pseudomonadales, Enterobacterales, and Bacteroidales MAGs, respectively; NRPS hybrids (Other NRPS) made up 16%, 15%, and 8% of BCGs annotated in Bacteroidales, Enterobacterales, and Pseudomonadales MAGs; and NRPS made up 22%, 9%, and 8% of BGCs annotated in Bacteroidales, Enterobacterales, and Pseudomonadales MAGs. Beta- lactone BCGs were found in Pseudomonadales (20%) and Enterobacterales (12%) but were absent in Bacteroidales. The only BGC class that was unique to Bacteroidales was hybrid T1PKS+hglE-KS (16%) (fig. 6C). MAGs from the genera *Gilliamella*, *Orbus*, *Entomomonas* possessed 23 biosynthetic gene clusters from 4 BGC classes: aryl polyene (n=2), NRPS (n=9), NRPS-like (n=2), NRPS+NRPS-like hybrid (n=2), and RiPP-like (n=8). The most abundant class of BGC in *Gilliamella*, *Orbus*, or *Entomomonas* MAGs were aryl polyene (n=2), RiPP-like (n=7), and NRPS (n=5), respectively (Fig. 7). Taken together, variation in the predicted BGCs among drosophilid-associated bacteria suggests that these bacterial can produce a diverse range of secondary biosynthetic molecules, and bacteria in different orders vary in the secondary metabolites they produce.

**Figure 6.**
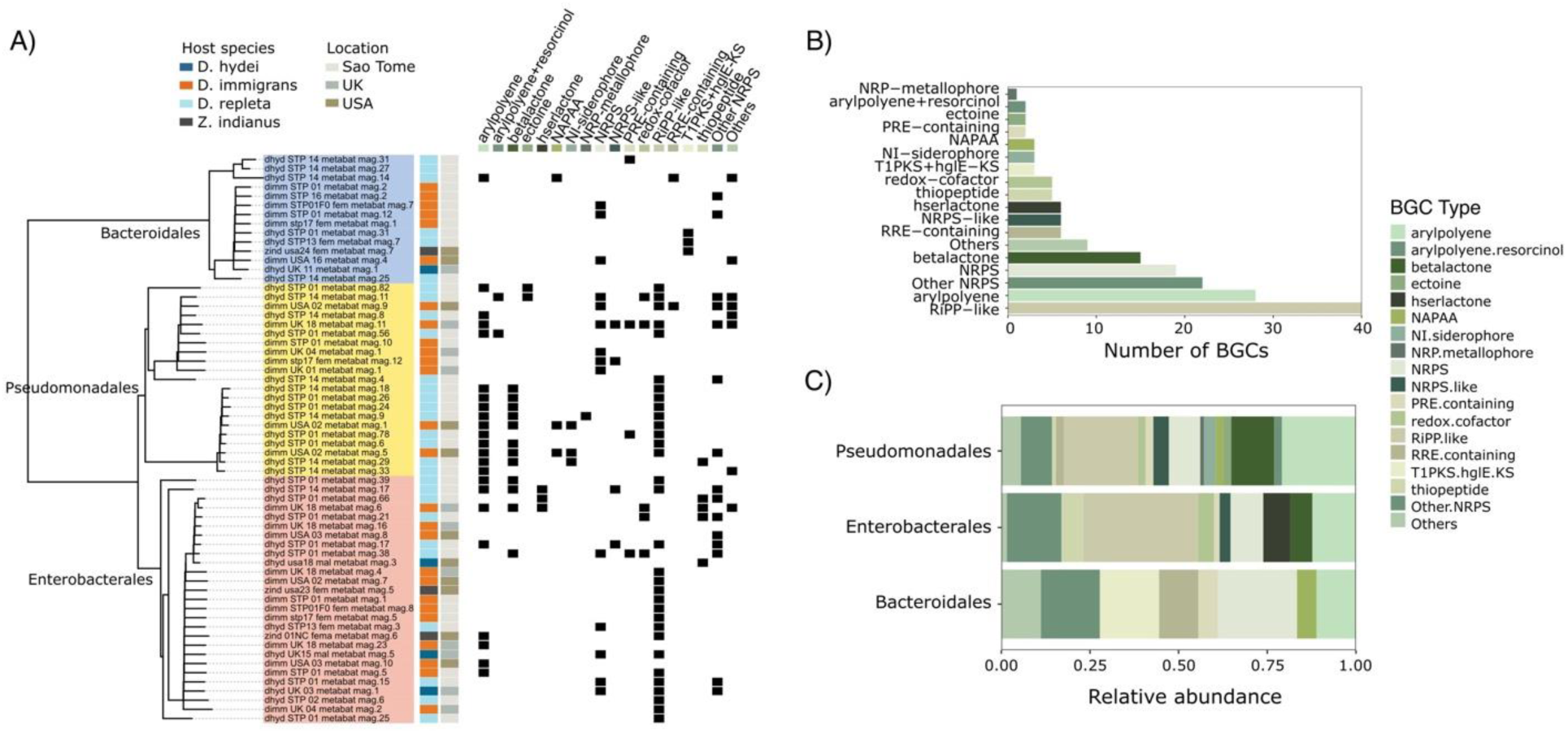
*Drosophila* MAGs differ in the secondary metabolite biosynthetic gene clusters (BGCs) they possess. **A**) Summary of biosynthetic products for MAGs from the Bacteriodales, Pseudomonadales, and Enterobacterales, including details of the host species and location the flies were collected from. Across MAGs, we identified 177 biosynthetic gene products belonging to 18 types of BGC (**B**). The relative abundance of BGC types was significantly different among the three focal orders of bacteria included in this analysis (**C**).

**Figure 7.**
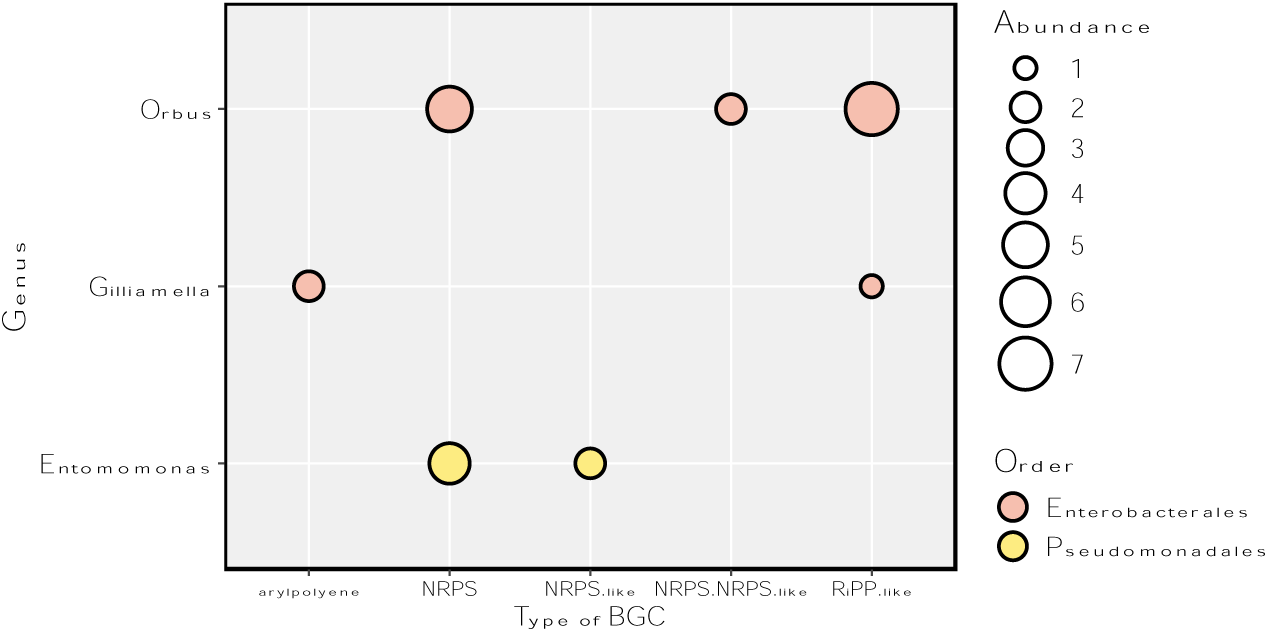
Abundance of biosynthetic gene clusters (BGCs) found in the classes (“Type of BGC”) that were the most common within the focal genera *Orbus*, *Gilliamella*, and *Entomomonas*.

## Discussion

Insect-associated bacteria can have diverse effects on their host’s biology—for example, they can detoxify dietary toxins (51) and provide protection from infection (7). However, there is also evidence that experimental manipulation of insect microbiomes (via antibiotic treatment) does not affect growth and development (52). These conflicting results highlight a need to identify the specific members of host-associated microbial communities and their functions. Consistent with previous studies of the microbiomes of wild drosophilids (10, 19, 20), we recovered a diverse set of host-associated bacterial reads from whole-organism sequencing (fig. 1). By assembling and analyzing MAGs from these reads, we found that the species of *Drosophila* (including *Zaprionus*) we study here are host to diverse lineages of bacteria (figs. 2-4) that differ in their functional gene content (figs. 5-7).

Bacteria in the family *Orbaceae* were among the most abundant taxa across samples, and we assembled MAGs from multiple host species and locations (figs 2-4). *Orbaceae* has previously been reported in metabarcoding studies of wild *Drosophila* as dominant members of their microbiome (19, 20) (19, 20, 53), and have also been reported from *Apis*, *Bombus*, and *Xylocopa* bees (25–27, 50, 54), *Eristalis* flys (55), and *Heliconius* and *Sasakia* butterflys (56, 57). These diverse hosts suggest that *Orbaceae* may be ‘core’ functional members of insect microbiomes. *Gilliamella apicola* is a species from the family *Orbaceae* found in bee hosts that has been shown to have functional metabolic capacities that complement other members of the honey bee gut microbiome (27), can detoxify toxic sugars found in the diets of bees (28), and can produce biosynthetic molecules that protect bees from infection by the bacterial pathogen *Melissococcus plutonius* (7). Using functional annotations, we found that *Drosophila*- associated *Orbaceae* MAGs are enriched for genes involved in pentose and glucuronate interconversions (map00040), ascorbate and aldarate metabolism (map00053), and resistance to beta-Lactam antibiotics (map01501) (fig. 4). Drosophilid- associated *Orbaceae* MAGs also harbored secondary metabolite biosynthetic gene clusters (BGCs) in the aryl polyene and RiPP-like classes. Ribosomally synthesized and post-translationally modified peptides (RiPP) have diverse roles, including playing a role in microbial interactions and antimicrobial activity (58, 59), and effects of RiPP-like BGCs on pathogens have been described from *Gilliamella* strains isolated bees (7). The diverse gene products produced by these bacteria highlight how insects and their microbiomes may be an important source of novel bioactive molecule discovery (60). While functional validation is needed—along with quantifying the impacts that drosophilid-associated *Orbaceae* have on host performance and fitness—, pathways and BSCs that are enriched in the drosophilid-associated MAGs we analyze here are candidates that could inform future functional studies. Indeed, a recently developed Pathfinder plasmid system has been verified in *Orbaceae* bacteria (61) and could be used to facilitate tests of candidate genes and pathways in *Orbaceae* isolated from diverse hosts.

In addition to bacteria from the family *Orbaceae*, we assembled four MAGs from the genus *Entomomonas* from *D. immigrans* collected in the USA and São Tomé and Príncipe (fig. 4C), and MAGs from the genus *Dysgonomonas* from *D. immigrans*, *D. repleta*, and *Z. indianus* collected from the USA and São Tomé and Príncipe (fig. 4B). Phylogenomic comparisons of drosophilid-associated *Entomomonas* MAGs to publicly available bacterial genomes showed that they are related to *Entomomonas* strains isolated from different insect orders (fig. 4C). By contrast, publicly available genomes from the genus *Dysgonomonas* are derived from both host-associated and environmental sources (fig. 4); however, *Dysgonomonas* has been reported as one of the core members of the microbiome of wild cactophilid *Drosophila* (53). By comparing relationships among drosophilid-associated MAGs and publicly available microbial genomes, our analyses suggest that *Orbaceae*, *Entomomonas*, and (to a lesser extent) *Dysgonomonas* are bacterial genera that may be evolved to utilize insects as hosts. As insect-associated microbial resources increase, identifying these ‘core’ members of the insect microbiome will facilitate tests of assembly and functional rules governing bacteria-insects interactions—for example, testing whether they represent generalist interactions or tightly coevolved symbioses (62).

We annotated an average of three BCGs for each drosophilid-associated MAG belonging to the Enterobacterales, Pseudomonadales, and Bacteriodales, indicating that members of the drosophilid microbiome have the capacity to produce potentially important secondary metabolites, despite generally not having large genomes (fig. S6). Secondary metabolites can play important roles in host-microbiome interactions and have previously been described in members of the gut microbiome of honey and bumble bees (7), herbivorous turtle ants (63) and mosquitoes (64). Secondary metabolites produced by BGCs can be involved in the regulation of symbiosis in fungus- faming termites (65), pathogenicity of malaria mosquitoes (64), and the deoxytication of β-Methylamino-L-alanine in cycad-feeding insects (66). BGCs belonging to the RiPP class were the most abundant in our drosophilid-associated MAGs, and may play a particularly important role in the microbiome since it has been observed that they can function to inhibit the growth of pathogens in bees (7) and can be involved in microbiome-host communication (67). Likewise, aryl polyenes (the second most abundant class in our dataset) have been observed to function as antioxidants, preventing stress caused by reactive oxygen species produced by the host in the case of bees (7, 68). Whether these molecules perform similar functions in drosophilid hosts remains to be confirmed; however, our results contribute to a growing body of work suggesting that BGCs possessed by the gut microbiome bacteria of insects contribute to the biology of their hosts and are a rich source of diverse secondary metabolites (7, 69).

*Wolbachia* and *Spiroplasma* are well known endosymbionts in insects, including in *Drosophila* (70–72), and we recovered sequences from these genera in four of our 31 samples. From these sequences were assembled MAGs identified as *Wolbachia* from both *Z. tsacasi* and *Z. taronus* hosts in our dataset. Both *Z. tsacasi* and *Z. taronus* are forest-dwelling species that we collected on the island of Saõ Tomé. *Wolbachia* infection frequencies have been shown to be highly variable among populations, species, and geographic regions (70, 73–76) and our results suggest that they are relatively rare across human-commensal drosophilids. Similarly, a screen of *Spiroplasma* in 35 species of *Drosophila* found that only three species—all from the “*repleta*” species group—were host to *Spiroplasma* infections (75). *Spiroplasma* infection rates in *D. hydei* (a member of the *repleta* species group) from the UK have been shown to vary from 15 to 29% across a 9-year period (77). The fact that we only found *Spiroplasma* in two samples of *D. hydei* from the UK is consistent with these past studies and may be indicative of a phylogenetic (and/or geographic) signal of *Spiroplasma* infection in *Drosophila* hosts. However, because we did not sample extensively in any one location, we are unable to estimate infection frequencies. Our data could prove useful for future phylogenetic or comparative genomic studies, and they provide novel MAGs from both *Spiroplasma* and *Wolbachia*.

Our results show that whole-organism reads generated using long-read ONT sequencers can be mined for metagenomic reads, and these can be used to estimate diversity and differences among host-associated microbial communities (fig. 1). ‘Mined’ bacterial reads can also be assembled into high-quality MAGs when sufficient bacterial sequence is extracted from the total sequence pool (fig. S2; table S2). Mining whole- organism reads to characterize host-associated microbes is likely to be a particularly useful approach when studying organisms where it is challenging to separate microbes from the host, or when microbial communities change when the organism is raised under artificial conditions. However, this approach is limited in not knowing where on the host the microorganisms are located. In many cases the location or life-history of the microorganisms can be reasonably inferred from knowledge of closely related taxa (e.g. endosymbionts and gut commensals); however, this information should be confirmed with additional species-specific data. Mining microbial reads from large datasets could be used to extract information from sequencing projects where characterizing the microbial community is not a primary goal. For example, recent work has used whole- organism sequencing of individual *Drosophila* spp. to generate genomic and phylogenetic resources for the group (78). This dataset includes wild-caught individuals whose data could be mined to quantify host-associated microbial diversity. Because laboratory and wild *Drosophila* show significant differences in the microbial communities they are host to (10, 19) accurate and transparent metadata need to be published alongside whole-organism sequencing to facilitate meaningful comparisons among host individuals. Moreover, our work highlights a need for functional studies of diverse insect microbiomes to gain a holistic view of the diversity, evolutionary history, and functional roles that insect-associated microorganisms play in their diverse hosts, and the ecosystems they inhabit.

## Acknowledgements

We thank Darren Obbard for sharing flies collected in the UK and Brandon Cooper for help collecting flies in São Tomé. This work was supported by a Royal Society Research Grant, The Royal Society of London (RGS\R1\221323) to AAC and the NERC Envision Doctoral Training Program (NE/L002604/1) awarded to AHO.

## Data Availability

Raw sequence reads are available on the NCBI SRA under BioProject PRJNA1188364: https://www.ncbi.nlm.nih.gov/bioproject/PRJNA1188364. Supporting figures, tables, scripts, and MAGs are available at Zenodo: doi:10.5281/zenodo.14173040: https://doi.org/10.5281/zenodo.14173040.

